# Modelling the Erythroblastic Island Niche of Dyserythropoietic Anaemia Type IV patients using Induced Pluripotent Stem Cells

**DOI:** 10.1101/2023.02.02.526657

**Authors:** Alisha May, Telma Ventura, Antonella Fidanza, Helena Volmer, Helen Taylor, Nicola Romanò, Sunita L D’Souza, James J. Bieker, Lesley M. Forrester

## Abstract

Congenital dyserythropoietic anaemia (CDA) type IV has been associated with an amino acid substitution, Glu325Lys (E325K), in the transcription factor KLF1. These patients present with a range of symptoms, including the persistence of nucleated red blood cells (RBCs) in the peripheral blood which reflects the known role for KLF1 within the erythroid cell lineage. The final stages of RBCs maturation and enucleation take place within the erythroblastic island (EBI) niche in close association with EBI macrophages. It is not known whether the detrimental effects of the E325K mutation in KLF1 are restricted to the erythroid lineage or whether deficiencies in macrophages associated with their niche also contribute to the disease pathology. To address this question, we generated an *in vitro* model of the human EBI niche using induced pluripotent stem cells (iPSCs) derived from a CDA type IV patient as well as iPSCs genetically modified to express an KLF1-E325K-ER^T2^ protein that could be activated with 4OH-tamoxifen. CDA patient-derived iPSCs and iPSCs expressing the activated KLF1-E325K-ER^T2^ protein showed significant deficiencies in the production of erythroid cells with associated disruption of some known KLF1 target genes. Macrophages could be generated from all iPSC lines but when the E325K-ER^T2^ fusion protein was activated, we noted the generation of a slightly less mature macrophage population marked by CD93. A subtle reduction in their ability to support RBC maturation was also associated with macrophages carrying the E325K-ER^T2^ transgene. Taken together these data support the notion that the clinically significant effects of the KLF1-E325K mutation are primarily associated with deficiencies in the erythroid lineage but that deficiencies in the niche might have the potential to exacerbate the condition. The strategy we describe provides a powerful approach to assess the effects of other mutations in KLF1 as well as other factors associated with the EBI niche.

## Introduction

The production of red blood cells (RBCs) at a rate of over two million per second is a complex and precisely controlled process that can be challenged by genetic deficiencies and alterations in environmental conditions. Severe, life-limiting anaemias resulting in reduced RBC numbers can be associated with congenital disease, chronic infection, inflammation and exposure to chemotherapeutic drugs (1, 2, 3, 4). The final steps of RBC maturation and enucleation occurs within the erythroblastic island (EBI) niche that consists of a central macrophage surrounded by developing erythroblasts (5, 6, 7, 8). EBI macrophages provide both positive and negative regulators of differentiation and development at various stages of erythroid maturation and have been associated with the pathological progression of some RBC disorders, including polycythemia vera and β-thalassemia (9, 10).

Congenital dyserythropoietic anaemia (CDA) type IV patients present with a range of symptoms, including nucleated RBCs in their peripheral blood, abnormalities in bone marrow erythroblasts, elevated fetal hemoglobin and iron overload (11, 12, 13, 14, 15, 16). One of the most severe forms of CDA type IV is associated with a single amino acid substitution Glu325Lys (E325K) in the second zinc finger of the erythroid specific transcription factor, KLF1. The mouse neonatal anaemia mutant (*Nan*) carries a semi-dominant mutation (E339D) in the equivalent DNA binding domain of the mouse protein and has been shown to reduce binding of DNA recognition sites and/or enable interaction with novel sites (17, 18). Expression profiling of erythroid cells from *Nan* mice and a CDA patient showed reduced expression of some known KLF1 target genes as well as ectopic expression of others (19, 20). These studies confirmed that this transcription factor is essential in the development and maturation of RBCs (21, 22, 23) but more recently KLF1 has also been associated with macrophages within the EBI niche (24, 25, 26).

Given the inaccessibility of the human EBI niche it has proven challenging to address whether deficiencies in EBI macrophages contribute to the disease pathology in CDA type IV patients that carry the E325K mutation. In this study we have used human induced pluripotent stem cells (iPSCs) to model CDA *in vitro* to confirm the intrinsic effects of the E325K mutation in erythroid cells and to assess the potential extrinsic effects of E325K in EBI-like macrophages. *In vitro* differentiation of CDA patient-derived iPSCs confirmed that the intrinsic effects of the E325K mutation in the erythroid lineage could be recapitulated using this strategy. We previously demonstrated that human EBI-like macrophages could be generated from iPSCs following 4OH-tamoxifen activation of a wild type KLF1-ER^T2^ fusion protein (25). Here we show that the presence of the E325K mutation alters this activity and has a subtle effect on the phenotype and function of iPSC-Derived Macrophages.

## Materials and Methods

### Establishment of patient-derived iPSC line

Induced pluripotent stem cell (iPSC) lines were derived from the patient’s blood mononuclear cells that were leftover from our previously published study that had received Institutional Review Board approval (12). No new patient material was obtained for the present study. After a brief culture to expand the erythroblast population a Sendai virus based approach (CytoTune 2.0 Kit, Invitrogen) was used to express the Yamanaka factors without any genomic integration (27). Clones were passaged >8x to dilute out the virus (verified by antibody staining), G-banded metaphase analysis showed a normal karyotype (not shown), and genomic DNA was sequenced to verify presence of the monoallelic E325K mutation (Supplementary Figure S1A).

### Human iPSC line maintenance

Human iPSC lines were maintained in StemProTM hESC SFM media (A1000701, Gibco) supplemented with 20 ng/ml human basic FGF (R&D) on either CELLstart Substrate (A1014201, Gibco) or Vitronectin (A31804, Gibco) coated wells. Media was changed daily. iPSCs were passaged when wells reached approximately 70-80% confluency with a StemPro EZPassage Disposable Stem Cell Passaging Tool (23181010, Gibco).

### Generation of pZDonor-AAVS1-CAG-HA-KLF1-E325K-ER^T2^-PolyA plasmid

The KLF1-E325K mutation (c973G>A) was introduced into an existing pZDonor-AAVS1-CAG-HA-KLF1-ER^T2^-PolyA plasmid (28) via site-directed mutagenesis using the Q5 Site-Directed Mutagenesis Kit (E0554S, New England BioLabs) following the manufacturer’s instructions. The forward KLF1 SDM NEB_FW and reverse KLF1 SDM NEB_RV primers were used (Supplementary Figure S2A and Supplementary Table S1).

### Transfection of iPSCs

SFCi55 iPSCs (28) were transfected using Xfect Transfection Reagent (631317, Takara Bio) following the manufacturer’s instructions. Puromycin selection was started at 2 μg/ml two days post transfection and increased to 4 μg/ml 4 days post transfection. Surviving colonies were picked approximately 2 weeks post transfection and grown into a 6-well plate format for screening.

### Erythroid differentiation of iPSCs

The protocol for the erythroid differentiation of iPSCs was adapted from Bernecker et al (29). Briefly, EBs are generated from iPSCs derived from fibroblasts with the addition of BMP4 (50 ng/ml), VEGF (50g/ml), and SCF (20ng/ml). EBs are plated in serum free differentiation media (30) supplemented with SCF (100 ng/ml), IL-3 (5 ng/ml) and EPO (3 U/ml) to generate haematopoietic cells. For KLF1 activation, 100 nM 4OH-tamoxifen (H6278-10MG Merck) was added every other day. Suspension cells were analysed at the point of harvest.

### Generation of human iPSC-derived macrophages

iPSC-derived macrophages were generated as previously described (25). For KLF1 activation, 100 nM 4OH-tamoxifen was added every other day.

### Erythroid differentiation of umbilical cord blood-derived CD34^+^ cells

Umbilical cord blood-derived CD34^+^ cells were cultured as previously described (25). For KLF1 activation, 100 nM 4OH-tamoxifen was added every other day.

### Cytospins

Cells for cytospins were suspended in PBS. Cells were cyto-centrifuged onto polysine slides at 500 rpm for 5 minutes in a Thermo Shandon Cytospin 4 and allowed to air-dry for 4-12 hours. Cells were fixed and stained using the Shandon™ Kwik-Diff™ Staining Kit (9990702, Thermo Fisher Scientific) following the manufacturer’s instructions.

### Immunohistochemistry

Cells were fixed into 96-well glass bottom plates (6055302, Perkin Elmer) in 4 % Formaldehyde (10231622, Fisher Scientific) for 15 minutes at room temperature. Cells were washed thrice with PBS and then permeabilised in PBS with 1 % BSA (A2153, Sigma-Aldrich) and 0.5 % Triton X-100 (X100, Sigma-Alrich) for 1 hour at room temperature. Cells were washed thrice with PBS before overnight incubation at 4°C in PBS with 1% donkey serum (ab7475, Abcam) and 1:200 Anti-EKLF/KLF1 antibody (ab2483, Abcam). Cells were then washed thrice with PBS and incubated for 1 hour at room temperature in PBS with 1 % donkey serum (ab7475, Abcam) and 1:1000 Donkey anti-Goat IgG (H+L) Cross-Absorbed Secondary Antibody, Alexa Fluor 647 (A-21447, Invitrogen). Cells were washed once with PBS and incubated with 1:1000 DAPI (D9542, Sigma-Aldrich) for 5 minutes at room temperature. Cells were washed twice with PBS and stored in PBS at 4 °C before imaging. Cells were imaged at 40X on the Opera Phenix® Plus High-Content Screening System and processed with Fiji software.

### Flow cytometry

Cells for analysis were resuspended in PBS with 1% BSA (A2153, Sigma-Aldrich) and 5 mM EDTA (15575020, Invitrogen). 1 × 10^5^ cells per sample were stained with appropriate antibodies for 15 minutes at room temperature. Samples were kept on ice until data collection using the LSR Fortessa (BD Biosciences) and BD FACSDIVA software. Data was analysed using FlowJo 10.8.1 software. Flow cytometry plots were gated using FMO controls. Briefly, single and live cells were gated, and then FMO controls were used to distinguish populations positive and negative for a specific marker. All antibodies used are listed in Supplementary Table S3.

### Gene expression analyses

RNA extraction was performed using the RNAeasy Mini Kit (74106, QIAGEN) following the manufacturer’s instructions. DNA was removed from samples using the RNase-free DNase Set (79254, QIAGEN). cDNA was generated from 500 ng of RNA per sample using the High-Capacity cDNA Reverse Transcription Kit (4368814, Thermo Fisher Scientific) following the manufacturer’s instructions. qRT-PCR reactions were performed on the Roche LightCycler® 480 Instrument. 2 ng of cDNA was amplified per reaction in a 364-well plate (4729749001, Roche) with LightCycler® 480 SYBR Green I Master (4887352001, Roche) according to the manufacturer’s instructions. All reactions were performed with 3 biological and 3 technical replicates. CT values were normalised to the reference gene GAPDH or the mean of the reference genes GAPDH and β-Actin. Data was analysed using the 2–ΔΔCt method. Graphs were generated and statistical analysis was performed using GraphPad Prism 8 software. Primers used are listed in SupplementaryTable S2.

### RNA-sequencing

RNA extraction was performed using the RNAeasy Mini Kit (QIAGEN) following the manufacturer’s instructions. DNA was removed from samples using the RNase-free DNase Set (QIAGEN). RNA quantity and quality was assessed using the Agilent 2100 Bioanalyser in conjuction with the RNA 6000 PicoLabChip Kit following manufacturer’s instructions. 35 automated TruSeq stranded mRNA-seq libraries from total RNA samples were generated by Edinburgh Genomics and sequenced using NovaSeq 100PE. Reads were trimmed using Cutadapt (version cutadapt-1.18-venv) (308). Reads were trimmed for quality at the 3’ end using a quality threshold of 30 for adaptor sequences of the TruSeq DNA kit (AGATCGGAAGAGC). Reads after trimming were required to have a minimum length of 50. The reference used for mapping was the Homo sapiens (GRCh38) genome from Ensembl. The annotation used for counting was the standard GTF-format annotation for that reference (annotation version 104). Reads were aligned to the reference genome using STAR (version 2.7.3a) specifying paired-end reads and the option --outSAMtype BAM Unsorted (309). All other parameters were left at default. Resulting BAM files were analysed using the DeSEQ2 package in R-4.2.1 for Windows. Genes were filtered and only genes with a count of 10 or higher in at least 2 samples were kept. Principal component analysis was undertaken on normalised and filtered expression data. For differential gene expression analyses, genes were filtered to include only genes with an adjusted p-value below 0.05.

## Results

### KLF1-E325K mutation in iPSCs affects erythroid differentiation

iPSCs were generated from peripheral blood mononuclear cells of a CDA Type IV patient and an unaffected individual (BM2.3) and confirmed the presence of the E325Kmutation in the genome of patient-derived iPSCs (Supplementary Figure S1A). These iPSC lines and a second control cell line (SFCi55), derived from skin fibroblasts (28), were differentiated into erythroid cells by adapting a previously published protocol (Figure 1A) (28, 29). All three cell lines generated a comparable percentage of CD43^+^ cells indicating that the E325K mutation did not affect commitment to the haematopoietic lineage (Figure 1B, C; Supplementary Figure 1B, C). There was no obvious effect on the proportion of cells expressing the early erythroid commitment marker, EpCAM. However, the percentage of cells expressing erythroid markers, CD235a (glycophorinA, GYPA) and CD71 (transferrin receptor, TRFC), that were generated from patient iPSCs was significantly lower compared to control iPSCs indicating a deficiency in erythropoietic differentiation (Figure 1B, C; Supplementary Figure 1B, C). Consistent with deficiency in erythroid cell production, a significant reduction in the expression of erythroid genes including *GYPA, TFRC, SLC4A1, HBA1* and *ICAM4* was observed (Figure 1D).

**Figure 1:**
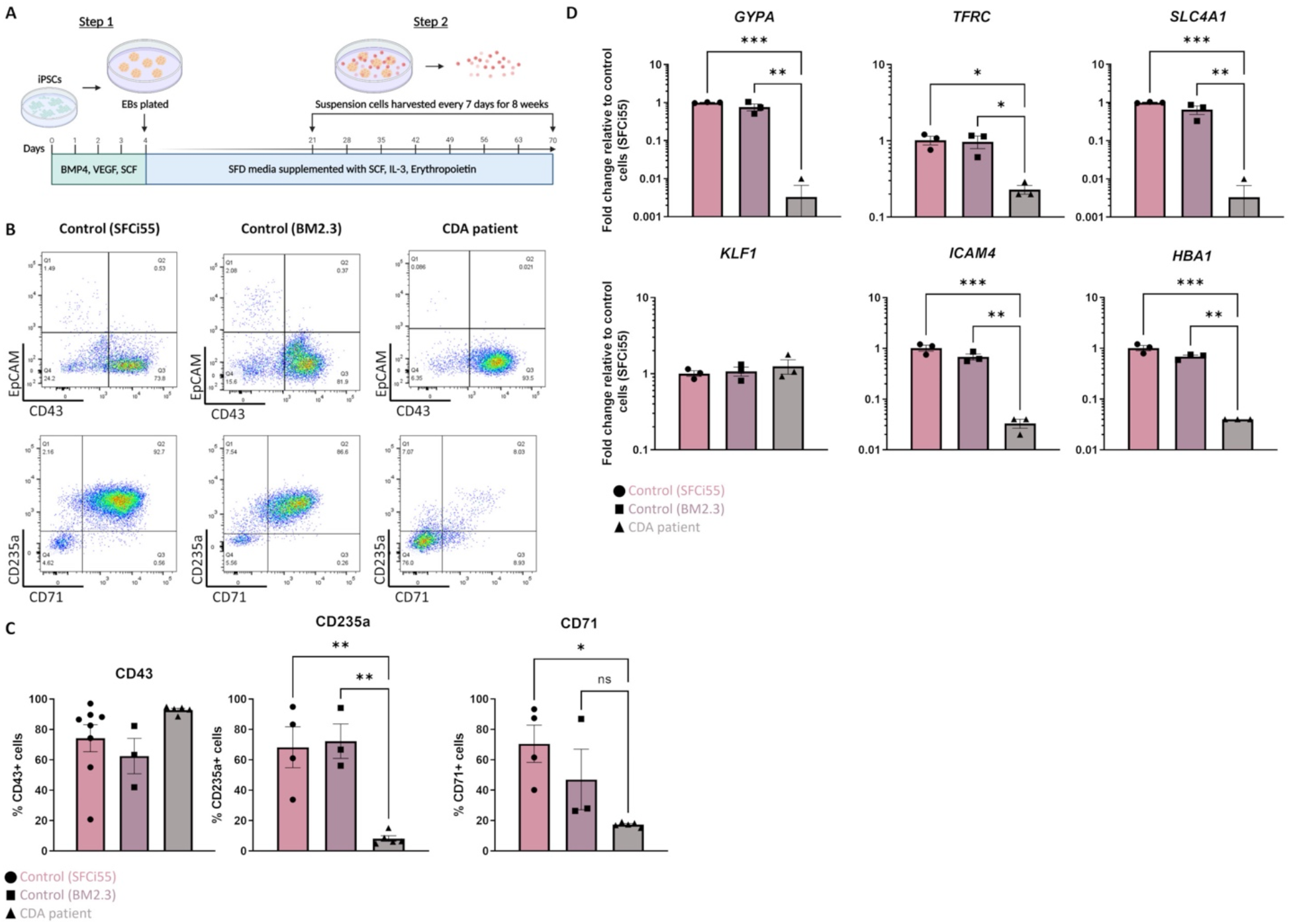
Erythroid progenitors generated from a CDA patient-iPSC line recapitulate disease pathology. A) Schematic of culture protocol to differentiate iPSC into erythroid cells. B) Representative flow cytometry plots of cell surface expression of CD43, EpCAM, CD235 and CD71 of suspension cells isolated from two control iPSC lines (SFCi55 and BM2.3) and one iPSC line derived from a CDA type IV patient. C) Quantification of flow cytometry analyses of cell surface marker expression. D) Gene expression analyses of *GYPA* (CD235a), *TRFC* (CD71), *SLC4A1* (*B*and 3), *HBA1* (hemoglobin subunit alpha), *ICAM4* and *KLF1* in suspension cells derived from two control and on CDA patient-derived iPSC line. Error bars represent SEM. One-way ANOVA with Tukey post-test. *p < 0.05, **p < 0.01

Given that we only had access to one iPSC line from a single patient and that isogenic controls were not available, we could not formally conclude that the erythroid differentiation deficiency was due to the E325K mutation and not associated with genetic background or simply a clonal iPSC effect. To address this concern, we generated an iPSC line where the KLF1-E325K protein could be activated by the addition of 4OH-tamoxifen (inducible/iKLF1-E325K) (28). We targeted the safe harbor *AAVS1* locus with a cassette carrying the *KLF1-E325K-ER*^*T2*^ fusion gene driven by the constitutively active CAG promoter and a puromycin resistance gene (Figure 2A) (31, 32). Following transfection of this targeting construct into parental SFCi55 iPSCs, puromycin-resistant clones were selected and screened by genomic PCR and Sanger sequencing to identify correctly targeted events (Supplementary Figure S2A,B). A targeting efficiency of 17% was achieved and two of the successfully targeted clones (named iCDA4.1 and iCDA4.20) were expanded and used in further experiments. Genomic PCR analyses using appropriate primers demonstrated that in the iCDA4.1 iPSC clone the targeting vector has integrated into only one of the *AAVS1* alleles whereas the iCDA4.20 was homozygous for the transgenic insertion and therefore had two copies of the transgene (Supplementary Figure S2A,B). We first tested whether the KLF1-E325K-ER^T2^ fusion protein would translocate to the nuclei in iPSCs upon addition of 4OH-tamoxifen using immunocytochemistry with an anti-KLF1 antibody (Figure 2B). In the absence of 4OH-tamoxifen, the majority of KLF1 staining was observed within the cytoplasm but upon addition of 4OH-tamoxifen, KLF1 was detected in the nucleus with a noticeable reduction in cytoplasmic staining. We noted comparable effects using the iKLF1.2 iPSC line that carried a wild type KLF1-ER^T2^ transgene (inducible/iKLF1-WT) previously reported indicating that the E325K mutation did not affect nuclear translocation of this fusion protein (28).

**Figure 2:**
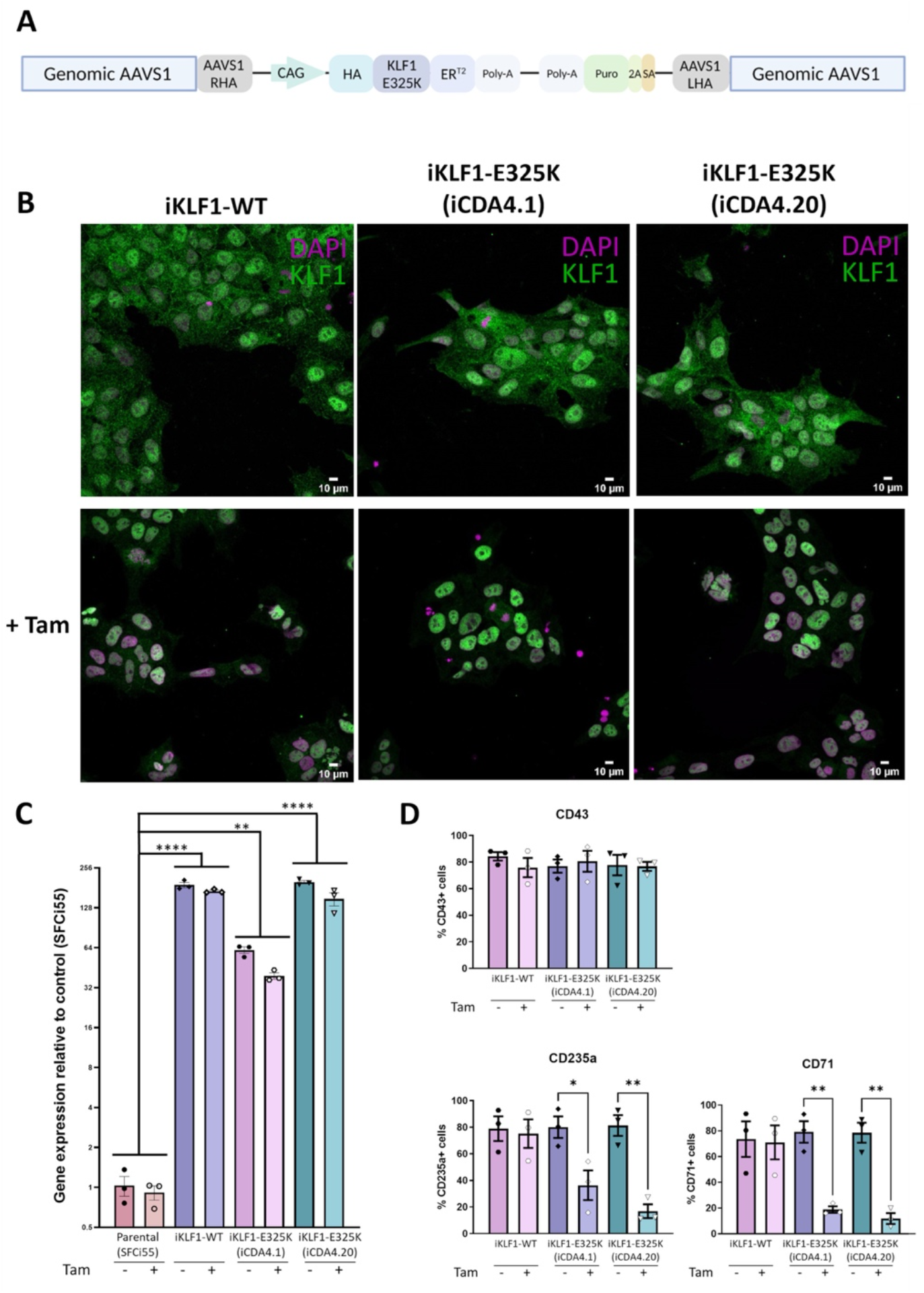
Inducible KLF1-E325K activation iPSC lines provide an alternative model for CDA. A) Schematic of the pZDonor-AAVS1-Puro-CAG-HA-KLF1-E325K-ERT2-PolyA construct integrated into the AAVS1 safe-harbor locus. B) Immunofluorescence staining of iPSCs from one inducible KLF1-WT (iKLF1-WT, iKLF1.2) and two inducible KLF1-E325K (iKLF1-E325K, iCDA4.1 and iCDA4.20) iPSC lines stained with an anti-KLF1 antibody (green) and the DAPI nuclear dye (magenta) in the presence (bottom panel) and absence (top panel) of 4OH-tamoxifen. 10uM scale bar. 40X magnification. C) Expression of KLF1 via qRT-PCR in RNA lysates taken from a iKLF1-WT and two iKLF1-E325K iPSC lines. Fold change relative to the parental control line (SFCi55). Error bars represent SEM. One-way ANOVA with Tukey post-test. *p < 0.05, **p < 0.01, ***p<0.0001. D) Quantification of flow cytometry analyses of cell surface expression of CD43, CD235a and CD71 of suspension cells harvested from erythroid differentiations of iKLF1-WT and iKLF1-E325K iPSCs. Error bars represent SEM. One-way ANOVA with Tukey post-test. *p < 0.05 **p < 0.01.

We previously reported that the level of endogenous KLF1 in control SFCi55 iPSCs was very low, so we next confirmed this and assessed the level of transgene expression by qRT-PCR (28). The level of KLF1 transcript detected by qRT-PCR was over 100-fold higher in the iPSCs carrying the CAG-driven KLF1-ER^T2^ and KLF1-E325K-ER^T2^ transgenes compared to control iPSCs (Figure 2C). The level of expression is significantly higher in the homozygous iCDA4.20 line compared to the heterozygous line iCDA4.1 line reflecting the number of copies of the transgene. The level of expression in the iCDA4.20 line was comparable to our previously reported iKLF1.2 iPSC line that also carried two copies of the transgene. As expected, the addition of 4OH-tamoxifen has no significant effect on the level of KLF1 transcript detected because 4OH-tamoxifen addition alters protein localization not *KLF1* transcriptional activity (Figure 2C).

We subjected the iKLF1.2, iCDA4.1 and iCDA4.20 iPSC lines to the same erythroid differentiation protocol that we used for the patient iPSC line (Figure 1A) (29). We first assessed the production of CD43^+^ haematopoietic progenitor cells (HPCs) in the presence and absence of 4OH-tamoxifen and noted that proportion of CD43 HPCs generated was comparable between all three cell lines and was unaffected by the addition of 4OH-tamoxifen (Figure 2D). In contrast, a significantly lower proportion of cells expressing the erythroid markers CD235a and CD71 were observed in the iCDA4.1 and iCDA4.20 cell lines following addition of 4OH-tamoxifen (Figure 2D). This 4OH-tamoxifen-induced reduction was not observed in the iKLF1.2 iPSC line nor in the control SFCi55 iPSC (Figure 2D; Supplementary Figure S2C). Notably, the effect of KLF1-E325K activation was more pronounced in the homozygous iCDA4.20 cell line, likely reflecting the higher level of expression of the KLF1-E325K transgene (Figure 2D). Taken together our *in vitro* iPSC-erythroid differentiation model system, at least at this level of analyses, recapitulates the CDA disease phenotype and is in keeping with the well-established role of KLF1 within the erythroid lineage.

### Activation of KLF1-E325K slightly impairs macrophage maturation

Macrophages can be differentiated from iPSCs and used to model the EI niche (25, 33, 34). Using these well-defined protocols, we generated macrophages from CDA patient iPSCs and demonstrated that they had a comparable cell surface phenotype to macrophages generated from control iPSCs and that their inclusion in culture could support the maturation of erythroid cells to the same extent (Supplementary Figure S3A-D). However, we previously demonstrated that the level of *KLF1* expression in iPSC-derived macrophages (iPSC-DMs) is very low but that increasing its activity using the *KLF1-ER*^*T2*^ transgene in iPSC-DMs resulted in a more EBI-like phenotype (25). We therefore hypothesized that increasing the levels of the mutant *KLF1-E325K* expression using our 4OH-tamoxifen-activatable system would be a more faithful model of the diseased EBI niche than macrophages derived from patient iPSCs.

To assess the effect of KLF1-E325K-ER^T2^ activation on macrophage production and phenotype, we carried out the macrophage differentiation protocol on the genetically manipulated iCDA4.1 and iCDA4.20 iPSC lines in the presence and absence of 4OH-tamoxifen. There was no obvious effect on the morphology of macrophages generated from each of these iPSC lines compared to control iKLF1-WT iPSC-DMs (Figure 3A). Almost all cells expressed macrophage markers (CD45, 25F9, CD169 and CD163) on the cell surface, and this was comparable in all cells when they were produced in either the presence or absence of 4OH-tamoxifen (Figure 3B). However, an interesting difference in the expression of the monocyte marker CD93 was observed (Figure 3C). The proportion of cells expressing CD93 was low (approximately <20%) in macrophages generated from control iKLF1-WT iPSCs and the addition of 4OH-tamoxifen reducing that to an even lower level (5%) (Figure 3C). In contrast, the proportion of cells expressing CD93 was significantly higher in macrophages that were differentiated from heterozygous iCDA4.1 iPSCs in the presence of 4OH-tamoxifen and an even higher proportion (40-60%) was observed in macrophages derived from the homozygous iCDA4.20 cell line that carried two copies of the transgene (Figure 3C). In both iKLF1-E325K lines, the proportion of CD93^+^ cells increased in macrophages treated with 4OH-tamoxifen. These data suggest that WT KLF1 expression drives macrophage maturation, the presence of the E325K mutation impairs that activity and that this phenotypic effect is associated to the level of expression of the mutant transgene (Figure 3C). The fact that a higher proportion of the immature, CD93-expressing cells was also noted in the iKLF1-E325K derived cells compared to controls in the absence of 4OH-tamoxifen likely reflects the fact that the ER domain is unable to entirely sequester all the molecules within the cytoplasmic region. This leakiness of the system was confirmed using immunocytochemistry on macrophages where KLF1 was detected in the nucleus of some cells in the absence of 4OH-tamoxifen (Supplementary Figure S3E).

**Figure 3:**
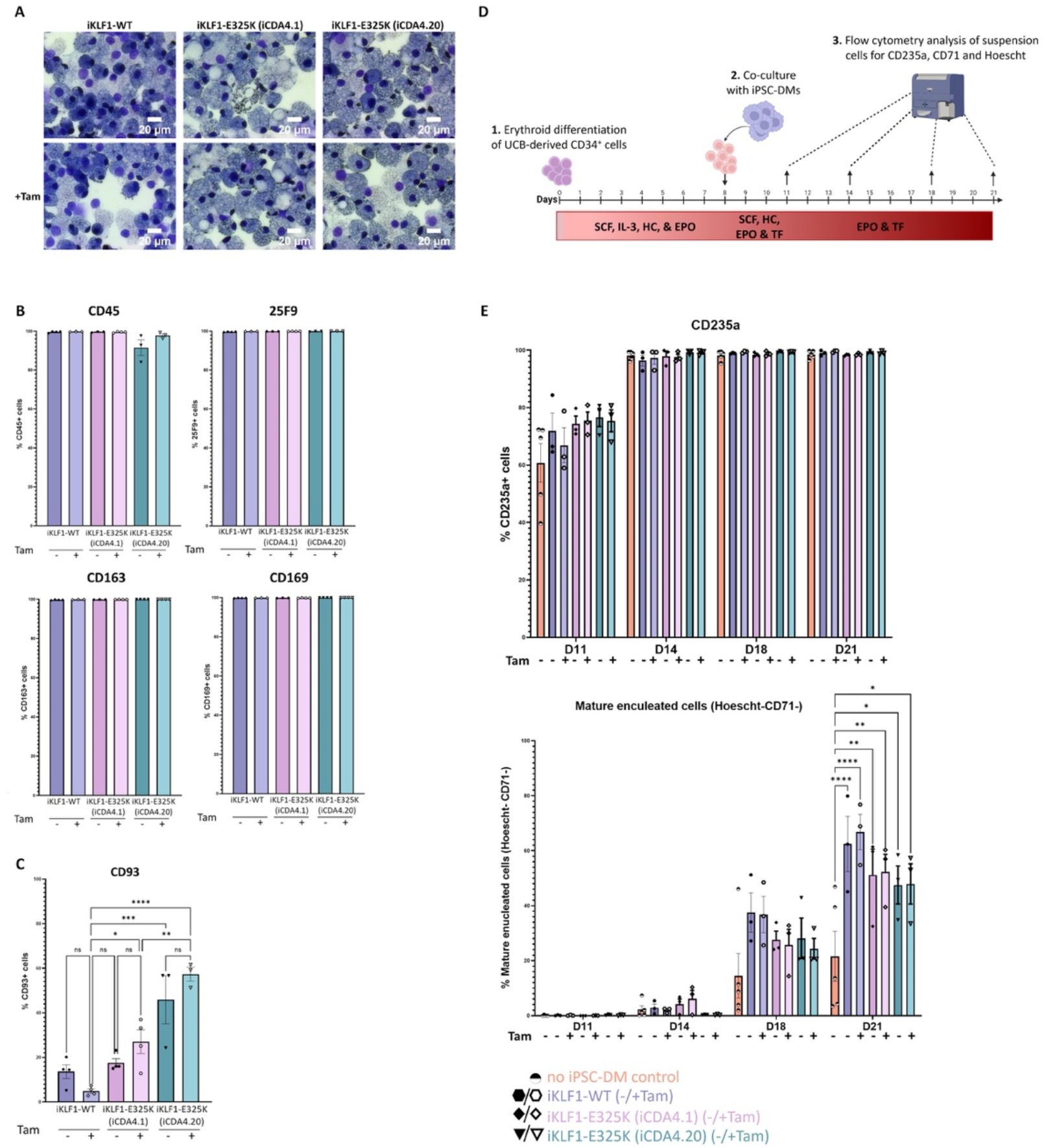
Activation of KLF1-E325K slightly impedes phenotypic and functional changes induced by KLF1. A) Representative KwikDiff stained cytospins of macrophages derived from a iKLF1-WT (iKLF1.2) and two iKLF1-E325K iPSC lines (iCDA4.1 and iCDA4.20). Flow cytometry analyses were performed on macrophages derived from a iKLF1-WT (iKLF1.2) and two iKLF1-E325K iPSC lines (iCDA4.1 and iCDA4.20) B) Quantification of flow cytometry analyses for cell surface marker expression of CD45, 25F9, CD163 and CD169. One-way ANOVA with Tukey post-test generated no statistically significant p-values. C) Quantification of flow cytometry analyses for cell surface marker expression of CD93. Error bars represent SEM. One-way ANOVA with Tukey post-test. *p < 0.05, **p < 0.01, ***p<0.0001. D) Schematic of *in vitro* erythroblastic island model protocol. Day 0-8: UCB-derived CD34+ cells are cultured with SCF, IL-3, hydrocortisone (HC) and EPO. Day 8-11: iPSC-DMs are added and cultured with the erythroid progenitors with SCF, HC, EPO and transferrin (TF). Day 11-21: Co-cultures are cultured with EPO and TF. At days 11, 14, 18 and 21 suspension cells are analysed by flow cytometry for expression of CD235a, CD71 and Hoechst. E) Quantification of flow cytometry analyses of suspension cells for CD235a, CD71 and Hoechst at days 11, 14, 18 and 21 of the culture. Error bars represent SEM. One-way ANOVA with Tukey post-test. *p < 0.05, **p < 0.01, ***p<0.0001

### Macrophages carrying the KLF1-E325K transgene are less efficient in supporting RBC maturation than those expressing WT-KLF1

We previously demonstrated that co-culture with iPSC-derived macrophages supported the erythroid differentiation and maturation of umbilical cord blood CD34^+^ HSPCs and that activation of KLF1 enhanced that activity (25). To address whether this enhancing effect is altered by the E325K mutation, we compared the phenotype of differentiating erythroid cells that were co-cultured with either iKLF1-WT or iKLF1-E325K iPSC-derived macrophages in an *in vitro* model of the EBI niche (Figure 3D). As previously described, erythroid maturation was assessed by flow cytometry using the cell surface marker CD71 that is lost as cells mature and the Hoeschst DNA stain that marks the presence of the nucleus in cells that have not undergone enucleation (25, 28). When no macrophages were present, the proportion of mature and enucleated erythroid cells (CD235a^+^/CD71^-^/Hoechst^-^) increased over the time course of the experiment to around 20% at day 21 (Figure 3E). Comparable to our previous reports, this was enhanced significantly to around 60% when co-cultured with iKLF1.2 macrophages with a slight further increase upon addition of 4OH-tamoxifen. Macrophages derived from iKLF1-E325K iPSCs (iCDA4.1and iCDA4.20) also enhanced the proportion of CD235a^+^/CD71^-^/Hoechst^-^ cells compared to cultures where no macrophages were present, but that increase was not as pronounced as the iKLF1-WT macrophages (Figure 3E). Activation of the E325K protein with 4OH-tamoxifen had no effect on that activity. Taken together these data suggest that the presence of the E325K mutation in iPSC-derived macrophages has an impact on their phenotype and function but the effect does not appear to have profound biological consequences in the specific context of supporting final erythroid differentiation.

### WT KLF1 has a more profound effect in the remodeling of macrophage transcriptome compared to KLF1 E325K

The E325K mutation in KLF1 is predicted to reduce binding to enhancers and promoters of target genes and/or enable interaction with novel DNA binding sites (11). We have used bulk RNA sequencing to assess how the presence of the E325K mutation affected the gene regulatory activity of KLF1 in iPSC-derived macrophages. We sequenced RNA derived from 5 replicates of macrophages derived from 3 iPSC lines (iKLF1.2, iCDA4.1 and SFCi55) in the presence and absence of 4OH-tamoxifen (30 samples in total). The parental SFCi55 cell line was used to identify any non-specific effects of 4OH-tamoxifen.

Differential gene expression analysis of macrophages derived from iKLF1-WT iPSCs identified 221 upregulated and 203 downregulated gene following 4OH-tamoxifen treatment (Figure 4A). Although this was significantly less than we had identified in our previous study, many of the genes were identified in both studies. In contrast, when assessing the effect of 4OH-tamoxifen treatment in macrophages derived from iKLF1-E325K iPSCs far fewer differentially expressed genes were identified; 17 genes were up-regulated and 2 were down-regulated in response to 4OH-tamoxifen (Figure 4A). Of the 221 genes that were up-regulated upon KLF1-activation, 5 were also up-regulated by mutant KLF1-E325K activation including *TRG-AS1, PHOSPHO1, SLC11A1* and *IL-33* (Figure 4B). These data suggested that activation of WT KLF1 in the iKLF1-WT macrophages had a greater effect on the transcriptome than the activation of mutant KLF1-E325K.

**Figure 4:**
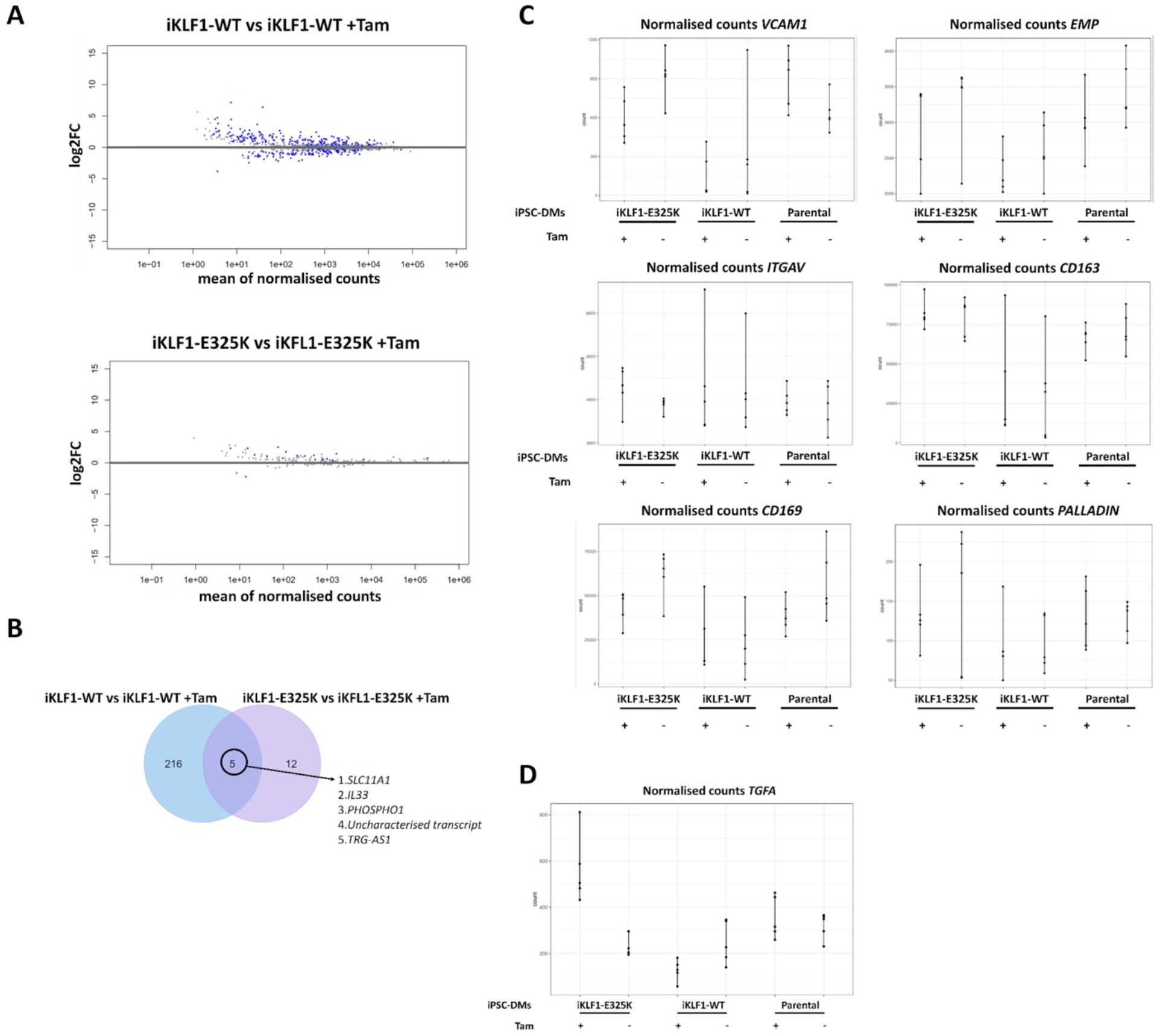
KLF1-E325K induces transcriptional changes in iPSC-DMs. A) Volcano plots for specified contrasts illustrating up-and down-regulated genes by Log2FC. Blue dots represent individual genes passing with an adjusted p-value below 0.05. C) Venn diagram of genes up-regulated in 4OH-tamoxifen treated iKLF1-WT iPSC-DMs and genes up-regulated in 4OH-tamoxifen treated iKLF1-E325K iPSC-DMs. D) Normalised counts shown for all 30 samples for the following genes encoding EBI macrophage attachment proteins: *VCAM1, EMP/MAEA, ITGAV, CD163, CD169* and *PALLADIN*. E) Normalised counts shown for all 30 samples for *TGFA*.

EBI macrophage attachment proteins were expressed in iPSC-derived macrophages and were unaffected by the presence of the E325K mutation (Figure 4C). The cell-cell contact of macrophages with erythroblasts within the EBI has been indicated to be more important to promoting erythroid cell maturation and enucleation than the secretion of factors (9, 25). We speculate that the predominantly retained ability of iKLF1-E325K iPSC-DMs to promote the maturation and enucleation of erythroid cells is due to cell-cell contact mediated by attachment proteins.

We identified TGFA to be the only gene that was significantly up-regulated upon KLF1-E325K activation but significantly down-regulated upon KLF1-WT activation (Figure 4D).

One of the most interesting group of genes were those that were upregulated by KLF1-WT but not KLF1-E325K because these are the most likely to be associated with any functional differences. We identified 4 genes that encoded secreted factors in this category, *ANGPTL7, ABI3BP, FDCSP*, and *IGFBP6*. IGFBP6 was particularly interesting, and *IGFBP6* expression was confirmed via qRT-PCR and was significantly upregulated in 4OH-tamoxifen treated iKLF1-WT but not iKLF1-E325K (iCDA4.1 and iCDA4.20) iPSC-DMs (Figure 5A). IGFBP6 was added in combination with NRG1, NOV, CCL13 and TNFSF10 to UCB-derived CD34^+^ cells under the erythroid cell differentiation conditions. Addition of these 5 factors increased the percentage of mature enucleated erythroid cells present in cultures at day 21, and removal of IGFBP6 resulted in a significant decrease in this population (unpublished data). We therefore wanted to investigate whether IGFBP6 alone promotes erythroid cell enucleation and maturation. The scaling up of protocols to generate RBCs for therapies need to be cost-effective, therefore identifying the key players in promoting RBC differentiations will enable a reduction in the numbers of costly cytokines that need to be added. Differentiating UCB-derived CD34^+^ cells were cultured alone or in the presence of two concentrations of IGFBP6, but we noted no significant effect on the percentage of CD235a^+^ cells nor on the percentage of mature enucleated erythroid cells (CD235a^+^/CD71^-^/Hoechst^-^) compared to the addition of no factors at all timepoints (Figure 5B).

**Figure 5:**
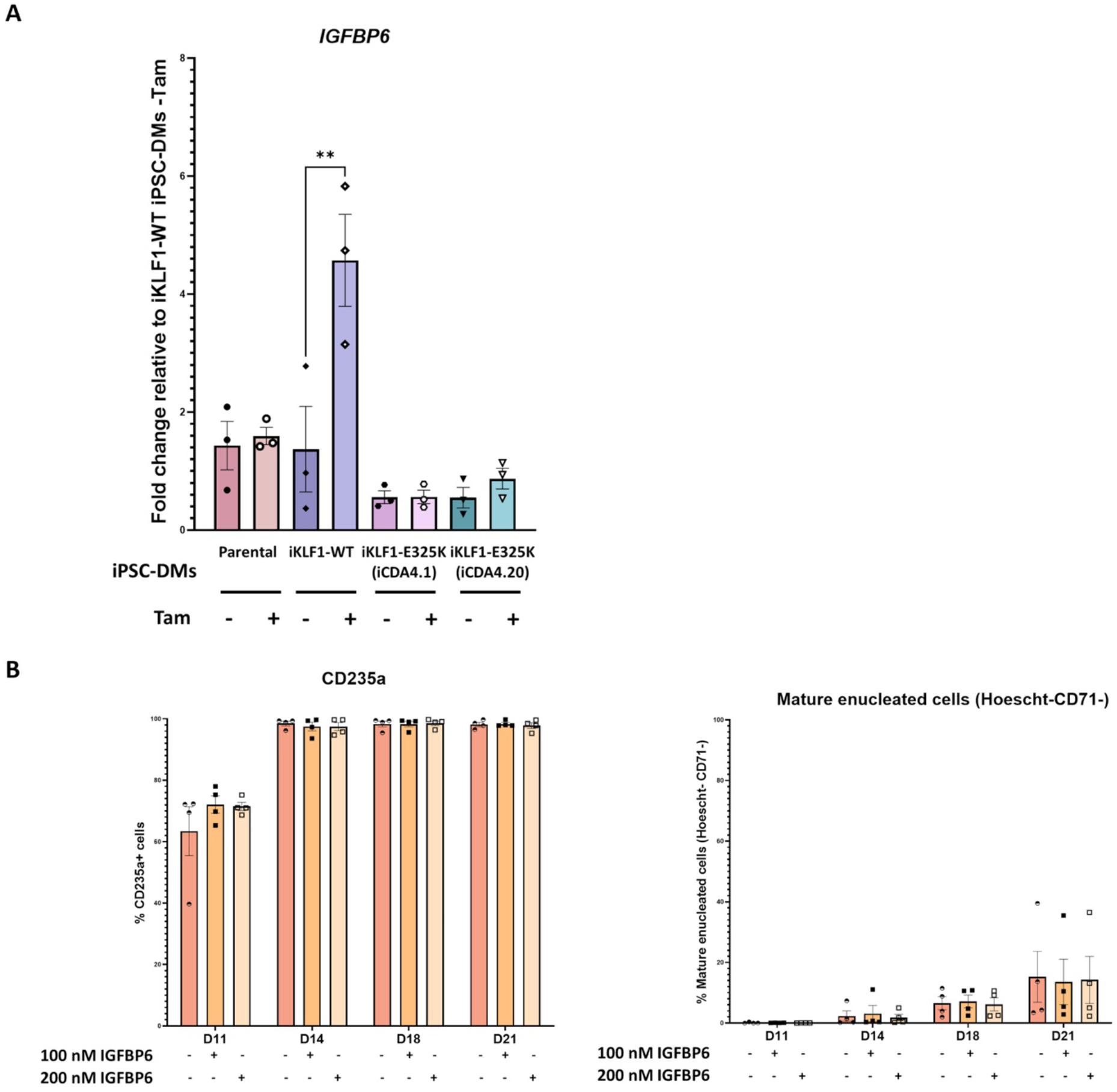
IGFBP6 alone has no effect on erythroid cell maturation and enucleation. A) Gene expression of IGFBP6 assessed by qRT-PCR. Error bars represent SEM. One-way ANOVA with Tukey post-test. **p < 0.01. B) IGFBP6 was added to UCB-derived CD34^+^ cells under erythroid differentiation conditions every 2 days at a concentration of either 100 nM or 200 nM. Quantification of flow cytometry analyses of suspension cells for CD235a, CD71 and Hoechst at days 11, 14, 18 and 21 of the culture. Error bars represent SEM. One-way ANOVA with Tukey post-test generated no statistically significant results.

## Discussion

Here we demonstrate that erythroid differentiation of CDA patient-derived iPSCs and iPSCs genetically modified to carry the E325K mutation in KLF1 provide a useful model to study erythroid deficiencies in CDA disease. We noted similar deficiencies in both models that were consistent with a previous report of an iPSC from a patient carrying the E325K mutation (35). That study showed no difference in the production of cells expressing CD34 and CD45 indicating that endothelial and haematopoietic commitment was unaffected by the presence of the E325K mutation. RNA sequencing of erythroid cells differentiated from CDA patient peripheral blood mononuclear cells *ex vivo* revealed the dysregulation of many erythroid genes and the ectopic expression of genes not normally expressed in erythroid cells (19). Consistent with this dysregulation, we observed a significant reduction in the expression of erythroid genes including *GYPA, TFRC, SLC4A1, HBA1* and *ICAM4* in erythroid differentiation of CDA patient-derived iPSCs.

We have previously shown that inducible activation of KLF1 in both hESCs and iPSCs enhances erythroid differentiation (28). Interestingly, we observed a more severe deficiency erythroid differentiations of the iPSC line that carried two copies of the E325K transgene compared with one copy. With the intrinsic effects of the E325K mutation in erythroid cells confirmed *in vitro*, we went on to assess the potential extrinsic effects of E325K in EBI-like macrophages.

Macrophages generated from CDA type IV patient iPSCs were phenotypically and functionally comparable to controls. This was not unexpected because iPSC-derived macrophages express low levels of KLF1 and so we predicted that patient iPSC-derived macrophages would be unlikely to accurately model the EBI macrophage. EBI macrophages have been demonstrated to express KLF1 (24, 26, 36), and we have previously shown that increased KLF1 expression and activation induces an EBI like phenotype in iPSC-derived macrophages (25).

Erythroid differentiation experiments demonstrated that the E325K inducible activation system could be used to recapitulate erythroid cell phenotype of CDA patient, so we therefore applied the system to investigate the effect of E325K on macrophages. Macrophages expressing E325K were morphologically comparable to controls and were predominantly positive for the cell surface markers CD45, 25F9, CD163 and CD169. However, we noted that fewer macrophages generated from the KLF1-E325K activation lines expressed CD93, and this was reduced upon KLF1-E325K activation. As iPSC-DMs mature, they lose expression of the monocyte marker CD93 (33). Activation of KLF1-WT reduced the percentage of CD93^+^ iPSC-DMs suggesting that KLF1-WT activation promotes iPSC-DM maturation. This supports our previous work showing that activation of KLF1-WT significantly increased the cell surface expression of the mature macrophage marker CD206 (25). As observed in erythroid differentiations, a more severe phenotypic effect was observed in the iPSC line that carried two copies of the E215K transgene. These data suggest that the E325K mutation impedes KLF1 induced macrophage maturation.

We also observed subtle differences in EBI niche support in the presence of E325K. Fewer erythroid cells matured and enucleated when co-cultured with iKLF1-E325K macrophages compared to iKLF1-WT macrophages. We previously reported a significant increase in the percentage of mature enucleated erythroid cells in cultures when KLF1-WT was activated in supporting macrophage, while in this study we only observe only a modest increase (25). We believe that this is likely due, in part, to the leakiness of the KLF1-ER^T2^ system.

It was anticipated that KLF1-WT would regulate more genes than KLF1-E325K for two reasons. Firstly, 4OH-tamoxifen treatment of iKLF1-WT iPSC-DMs activates KLF1-WT, the fully functional protein that has been previously identified to regulate gene expression in EBI macrophages (24, 25). Secondly, iKLF1-WT iPSCs have two copies of the KLF1-WT-ER^T2^ transgene while iKLF1-E325K iPSCs have only one copy of the iKLF1-E325K-ER^T2^ transgene. Indeed, RNA-sequencing indicates that there far fewer genes are regulated by the mutant E325K protein compared to the wild type KLF1 protein.

While we observed a reduced number of mature and enucleated erythroid cells in *in vitro* EBI cultures with KLF1-E325K macrophages compared to the KLF1-WT control, KLF1-E325K retained the ability to support the maturation and enucleation of a significant percentage of cells. We speculate that this retained function within the EBI is due to EBI macrophage protein interactions. The association of erythroblasts with macrophages promotes erythroid cell maturation and enucleation, and elimination of this contact using a transwell has been demonstrated to significantly decrease erythroid cell maturation and enucleation (9, 25). We found that expression of the following genes encoding EBI attachment proteins were unaltered in iCDA4.1 macrophages: *VCAM1, EMP/MAEA, ITGAV, CD163, CD169* and *PALLADIN*.

We previously identified up-regulation of the genes encoding the secreted factors IL-33, SERPINB2, and ANGPTL7 upon activation of KLF1-WT in iPSC-derived macrophages (25). Addition of these secreted factors to erythroid differentiations of UCB CD34^+^ cells significantly increased the percentage of mature enucleated cells in the cultures. Removal of any individual factor resulted in a significant reduction in mature enucleated cells, with removal of IL-33 resulting in the most significant reduction. Addition of IL-33 alone did not increase erythroid cell maturation and enucleation, indicating that IL-33 acts in synergy with other factors. In agreement with this previous study, IL-33 and ANGPTL7 were up-regulated in iPSC-DMs upon KLF1-WT activation in our dataset. SERPINB2 was not up regulated and was not identified to be expressed in any macrophages in our dataset. In addition to the leakiness of our system, this lack of SERPINB2 expression in could be responsible for the modest increase in mature enucleated cells we observed in cultures with KLF1-WT activated macrophages. In contrast to KLF1-WT activation, KLF1-E325K activation up-regulated IL-33 but not ANGPTL7. This loss of ANPTL7 expression could explain this reduction in the numbers of mature enucleated erythroid cells, as IL-33 has been previously shown to promote erythroid cell maturation and enucleation only in combination with ANPTL7 and/or SERPINB2

We speculated that loss of IGFBP6 regulation by E325K could explain the subtle defects in erythroid maturation and enucleation observed in cultures with E325K compared to KLF1-WT macrophages. To test this hypothesis, we added IGFBP6 to our erythroid differentiation of UCB CD34^+^ cells but observed no differences compared to control cultures. While IGFBP6 alone has no effect, we previously identified a decrease in erythroid cell maturation and enucleation when IGFBP6 was excluded from a cocktail of added secreted factors. These data suggest that, similarly to IL-33, IGFBP6 acts in association with other secreted factors to promote erythropoiesis.

TGFA was the only gene that was up-regulated by KLF1-E325K activation and down-regulated by KLF1-WT activation. TGFA encodes for the protein transforming growth factor alpha (TGF-α), a member of the epidermal growth factor which activates a signaling pathway for cell proliferation (37). In macrophages, TGF-α expression and secretion has only been reported by alveolar macrophages (38, 39). TGF-α addition to avian erythroid progenitors was shown to promote their self-renewal, and removal of TGF-α from erythroid progenitors from chick bone marrow caused them to terminally differentiate (40, 41, 42). It is important to note that avian erythroid cells do not enucleate as part of terminal erythroid differentiation, while human erythroid cells do. Thus, the presence of a higher TGFA concentration within the EBI niche might also prevent erythroid cells from fully maturing. Further experiments are needed to investigate whether TGF-α also promotes the self-renewal of human erythroid progenitors, and what if any effect this has on cell cycle exit, which is required for erythroblast enucleation (22).

## Conclusion

Collectively this study demonstrates that genetically modified iPSCs provide an *in vitro* model to study mechanisms associated RBC disorders both within the erythroid lineage itself as well as in cells associated with the erythroblastic island niche. This strategy could be used to model the EBI niche of other RBC disorders to assess the possible contributions of EBI macrophages and to potentially discover new druggable targets. Existing treatments for several hereditary and acquired RBC disorders are effective in managing some of these conditions, but few offer long term cures. Finding new treatments relies on the full understanding of the cellular and molecular interactions associated with the production and maturation of RBCs within the EBI niche. This strategy would prove especially useful for rare diseases such as CDA type IV for which there is very limited availability of primary cells, and in which animal models do not exactly recapitulate the disease.

## Acknowledgements

This work was supported by the Wellcome Trust [108906/Z/15/Z](AM); College of Medicine and Veterinary Medicine,University of Edinburgh (AM), UKRI Medical Research Council [MR/ T013923/1](LF, TV), UKRI Biotechnology and Biological Sciences Research Council [BB/S002219/1](AF, HT) and a US PHS award [DK121671](JJB). We thank Fiona Rossi (CRM flow cytometry facility), Mattieu Vermenen and Justyna Cholewa-Waclaw (CRM imaging facility) and Leslie Nitsche for her help with RNA sequence analyses. The data described in the manuscript will also be available online as AM’s PhD thesis University of Edinburgh.

## References

1. Gomes AC, Gomes MS. Hematopoietic niches, erythropoiesis and anemia of chronic infection. Experimental hematology. 2016;44(2):85–91.

2. Iskander D, Psaila B, Gerrard G, Chaidos A, En Foong H, Harrington Y, et al. Elucidation of the EP defect in Diamond-Blackfan anemia by characterization and prospective isolation of human EPs. Blood. 2015;125(16):2553–7.

3. Perkins A, Xu X, Higgs DR, Patrinos GP, Arnaud L, Bieker JJ, et al. “Kruppeling” erythropoiesis: an unexpected broad spectrum of human red blood cell disorders due to KLF1 variants unveiled by genomic sequencing. Blood. 2016;127(15):1856–62.

4. J.T. P, X.T. G. Erythropoiesis - genetic abnormalities. In: Elliott SG, Foote MA, Molineux G, editors. Erythropoietins, Erythropoietic Factors and Erythropoiesis Milestones in Drug Therapy. Basel, Switzerland: Birkhäuser Verlag; 2009.

5. Manwani D, Bieker JJ. The erythroblastic island. Curr Top Dev Biol. 2008;82:23–53.

6. Ji P, Murata-Hori M, Lodish HF. Formation of mammalian erythrocytes: chromatin condensation and enucleation. Trends Cell Biol. 2011;21(7):409–15.

7. Bessis M. [Erythroblastic island, functional unity of bone marrow]. Rev Hematol. 1958;13(1):8–11.

8. Chasis JA, Mohandas N. Erythroblastic islands: niches for erythropoiesis. Blood. 2008;112(3):470–8.

9. Chow A, Huggins M, Ahmed J, Hashimoto D, Lucas D, Kunisaki Y, et al. CD169(+) macrophages provide a niche promoting erythropoiesis under homeostasis and stress. Nat Med. 2013;19(4):429–36.

10. Ramos P, Casu C, Gardenghi S, Breda L, Crielaard BJ, Guy E, et al. Macrophages support pathological erythropoiesis in polycythemia vera and beta-thalassemia. Nat Med. 2013;19(4):437–45.

11. Arnaud L, Saison C, Helias V, Lucien N, Steschenko D, Giarratana MC, et al. A dominant mutation in the gene encoding the erythroid transcription factor KLF1 causes a congenital dyserythropoietic anemia. Am J Hum Genet. 2010;87(5):721–7.

12. Jaffray JA, Mitchell WB, Gnanapragasam MN, Seshan SV, Guo X, Westhoff CM, et al. Erythroid transcription factor EKLF/KLF1 mutation causing congenital dyserythropoietic anemia type IV in a patient of Taiwanese origin: review of all reported cases and development of a clinical diagnostic paradigm. Blood Cells Mol Dis. 2013;51(2):71–5.

13. Wickramasinghe SN, Illum N, Wimberley PD. Congenital dyserythropoietic anaemia with novel intra-erythroblastic and intra-erythrocytic inclusions. Br J Haematol. 1991;79(2):322–30.

14. Singleton BK, Fairweather VSS, Lau W, Parsons SF, Burton NM, Frayne J, et al. A Novel EKLF Mutation in a Patient with Dyserythropoietic Anemia: The First Association of EKLF with Disease in Man. Blood. 2009;114(22):162-.

15. Ortolano R, Forouhar M, Warwick A, Harper D. A Case of Congenital Dyserythropoeitic Anemia Type IV Caused by E325K Mutation in Erythroid Transcription Factor KLF1. J Pediatr Hematol Oncol. 2018;40(6):e389–e91.

16. de-la-Iglesia-Iñigo S, Moreno-Carralero MI, Lemes-Castellano A, Molero-Labarta T, Méndez M, Morán-Jiménez MJ. A case of congenital dyserythropoietic anemia type IV. Clin Case Rep. 52017. p. 248–52.

17. Siatecka M, Sahr KE, Andersen SG, Mezei M, Bieker JJ, Peters LL. Severe anemia in the Nan mutant mouse caused by sequence-selective disruption of erythroid Kruppel-like factor. Proc Natl Acad Sci U S A. 2010;107(34):15151–6.

18. Planutis A, Xue L, Trainor CD, Dangeti M, Gillinder K, Siatecka M, et al. Neomorphic effects of the neonatal anemia (Nan-Eklf) mutation contribute to deficits throughout development. Development. 2017;144(3):430–40.

19. Varricchio L, Planutis A, Manwani D, Jaffray J, Mitchell WB, Migliaccio AR, et al. Genetic disarray follows mutant KLF1-E325K expression in a congenital dyserythropoietic anemia patient. Haematologica. 2019;104(12):2372–80.

20. Nébor D, Graber JH, Ciciotte SL, Robledo RF, Papoin J, Hartman E, et al. Mutant KLF1 in Adult Anemic Nan Mice Leads to Profound Transcriptome Changes and Disordered Erythropoiesis. Sci Rep. 2018;8(1):12793.

21. Siatecka M, Bieker JJ. The multifunctional role of EKLF/KLF1 during erythropoiesis. Blood. 2011;118(8):2044–54.

22. Gnanapragasam MN, McGrath KE, Catherman S, Xue L, Palis J, Bieker JJ. EKLF/KLF1-regulated cell cycle exit is essential for erythroblast enucleation. Blood. 2016;128(12):1631–41.

23. Tallack MR, Whitington T, Yuen WS, Wainwright EN, Keys JR, Gardiner BB, et al. A global role for KLF1 in erythropoiesis revealed by ChIP-seq in primary erythroid cells. Genome Res. 2010;20(8):1052–63.

24. Xue L, Galdass M, Gnanapragasam MN, Manwani D, Bieker JJ. Extrinsic and intrinsic control by EKLF (KLF1) within a specialized erythroid niche. Development. 2014;141(11):2245–54.

25. Lopez-Yrigoyen M, Yang CT, Fidanza A, Cassetta L, Taylor AH, McCahill A, et al. Genetic programming of macrophages generates an in vitro model for the human erythroid island niche. Nat Commun. 2019;10(1):881.

26. Mukherjee K, Xue L, Planutis A, Gnanapragasam MN, Chess A, Bieker JJ. EKLF/KLF1 expression defines a unique macrophage subset during mouse erythropoiesis. Elife. 2021;10.

27. Carcamo-Orive I, Hoffman GE, Cundiff P, Beckmann ND, D’Souza SL, Knowles JW, et al. Analysis of Transcriptional Variability in a Large Human iPSC Library Reveals Genetic and Non-genetic Determinants of Heterogeneity. Cell Stem Cell. 2017;20(4):518-32.e9.

28. Yang CT, Ma R, Axton RA, Jackson M, Taylor AH, Fidanza A, et al. Activation of KLF1 Enhances the Differentiation and Maturation of Red Blood Cells from Human Pluripotent Stem Cells. Stem Cells. 2017;35(4):886–97.

29. Bernecker C, Ackermann M, Lachmann N, Rohrhofer L, Zaehres H, Araúzo-Bravo MJ, et al. Enhanced Ex Vivo Generation of Erythroid Cells from Human Induced Pluripotent Stem Cells in a Simplified Cell Culture System with Low Cytokine Support. Stem Cells Dev. 2019;28(23):1540–51.

30. Sturgeon CM, Chicha L, Ditadi A, Zhou Q, McGrath KE, Palis J, et al. Primitive erythropoiesis is regulated by miR-126 via nonhematopoietic Vcam-1+ cells. Dev Cell. 2012;23(1):45–57.

31. Smith JR, Maguire S, Davis LA, Alexander M, Yang F, Chandran S, et al. Robust, persistent transgene expression in human embryonic stem cells is achieved with AAVS1-targeted integration. Stem Cells. 2008;26(2):496–504.

32. Hockemeyer D, Soldner F, Beard C, Gao Q, Mitalipova M, DeKelver RC, et al. Efficient targeting of expressed and silent genes in human ESCs and iPSCs using zinc-finger nucleases. Nat Biotechnol. 2009;27(9):851–7.

33. Lopez-Yrigoyen M, May A, Ventura T, Taylor H, Fidanza A, Cassetta L, et al. Production and Characterization of Human Macrophages from Pluripotent Stem Cells. J Vis Exp. 2020(158).

34. May A, Forrester LM. The erythroblastic island niche: modeling in health, stress, and disease. Exp Hematol. 2020;91:10–21.

35. Kohara H, Utsugisawa T, Sakamoto C, Hirose L, Ogawa Y, Ogura H, et al. KLF1 mutation E325K induces cell cycle arrest in erythroid cells differentiated from congenital dyserythropoietic anemia patient-specific induced pluripotent stem cells. Exp Hematol. 2019;73:25-37.e8.

36. Mukherjee K, Bieker JJ. Transcriptional Control of Gene Expression and the Heterogeneous Cellular Identity of Erythroblastic Island Macrophages. Front Genet. 2021;12:756028.

37. Coffey RJ, Gangarosa LM, Damstrup L, Dempsey PJ. Basic actions of transforming growth factor-alpha and related peptides. Eur J Gastroenterol Hepatol. 1995;7(10):923–7.

38. Madtes DK, Raines EW, Sakariassen KS, Assoian RK, Sporn MB, Bell GI, et al. Induction of transforming growth factor-alpha in activated human alveolar macrophages. Cell. 1988;53(2):285–93.

39. Wagner CL, Ryan RM, Forsythe DE, Keenan A, Finkelstein JN. Secretion of transforming growth factor-alpha (TGF alpha) by postnatal rabbit alveolar macrophages. Pediatr Res. 1995;38(1):49–54.

40. Hayman MJ, Meyer S, Martin F, Steinlein P, Beug H. Self-renewal and differentiation of normal avian erythroid progenitor cells: regulatory roles of the TGF alpha/c-ErbB and SCF/c-kit receptors. Cell. 1993;74(1):157–69.

41. Gandrillon O, Schmidt U, Beug H, Samarut J. TGF-beta cooperates with TGF-alpha to induce the self-renewal of normal erythrocytic progenitors: evidence for an autocrine mechanism. Embo j. 1999;18(10):2764–81.

42. Schroeder C, Gibson L, Nordström C, Beug H. The estrogen receptor cooperates with the TGF alpha receptor (c-erbB) in regulation of chicken erythroid progenitor self-renewal. Embo j. 1993;12(3):951–60.

