## Supplementary Files and Tables for "Modelling the Erythroblastic Island Niche of Dyserythropoietic Anaemia Type IV patients using Induced Pluripotent Stem Cells"

**May et al. Supplementary Figures and Tables**

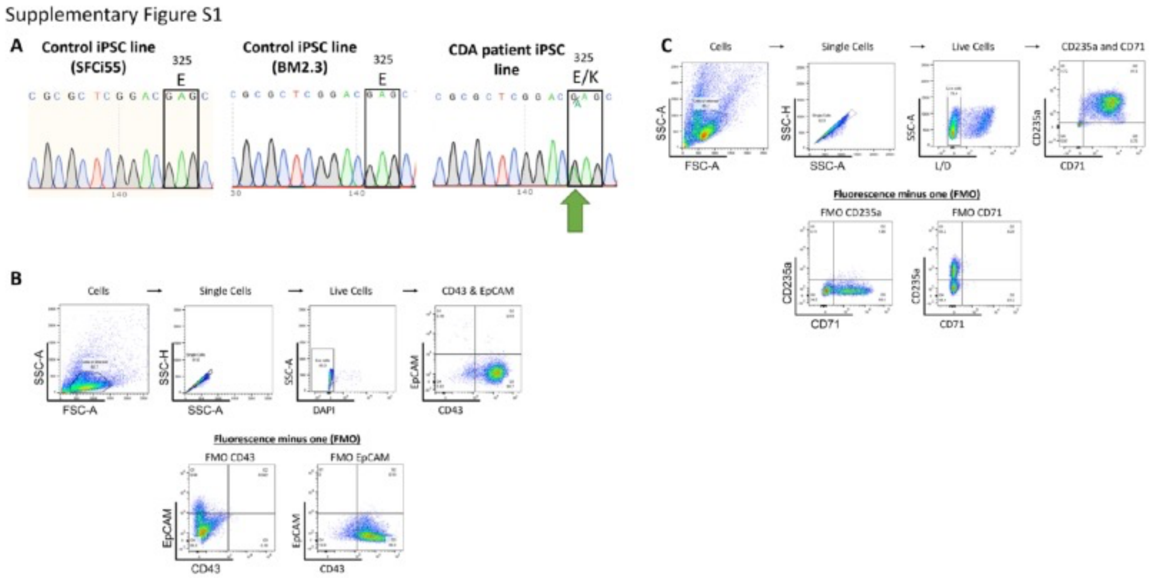


**Supplementary Figure S1:**
A) Sequence traces showing the WT c.973G locus in two KLF1-WT iPSC lines (SFCi55 and BM2.3) and the mutant c.973G>A locus CDA type IV patient-iSPC-derived line. B) Gating strategy for analyses of suspension cells harvested from erythroid differentiations. Single, live cells were gated and analysed for expression of CD43 and EpCAM using fluorescence minus one (FMO) to set gates. C) Gating strategy for analyses of suspension cells harvested from erythroid differentiations. Single, live cells were gated and analysed for expression of CD235a and CD71 using fluorescence minus one (FMO) to set gates.

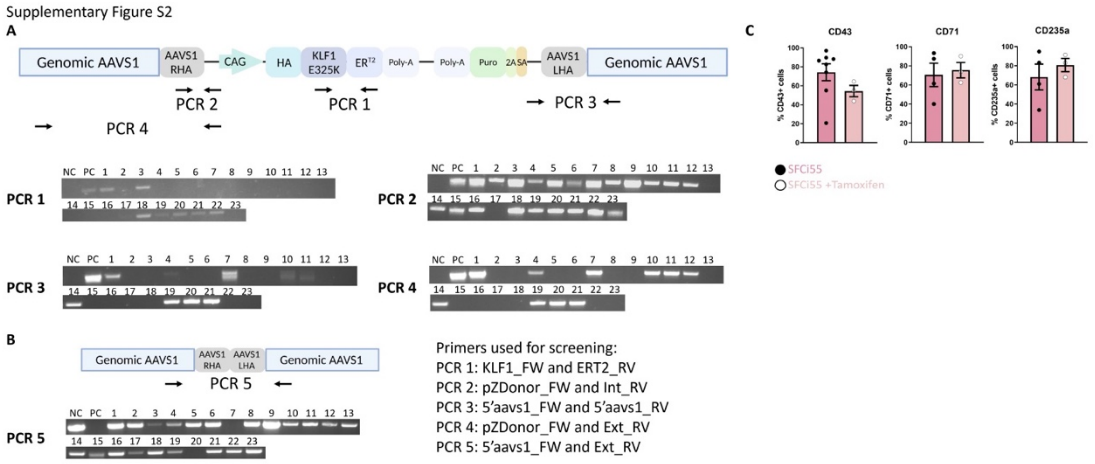


**Supplementary Figure S2:**

A) A schematic diagram of the KLF1-E325K-ER$^{T2}$ construct targeted into the AAVS1 locus showing the amplicons for PCRs 1-4 and the PCR screening results for clones 1-23. Negative control (NC) is SFCi55 iPSCs. Positive control (PC) is inducible KLF1-WT iPSCs. B: A schematic diagram of the non-targeted AAVS1 locus showing the amplicon for PCR 5 and the PCR screening results for the NC, PC and clones 1-23 C) Quantification of flow cytometry analyses of the percentage of cells generated from erythroid differentiations from parental line iPSCs (SFCi55) expressing CD43, CD71 and CD235a. Unpaired t-tests generated no statistically significant p-values.

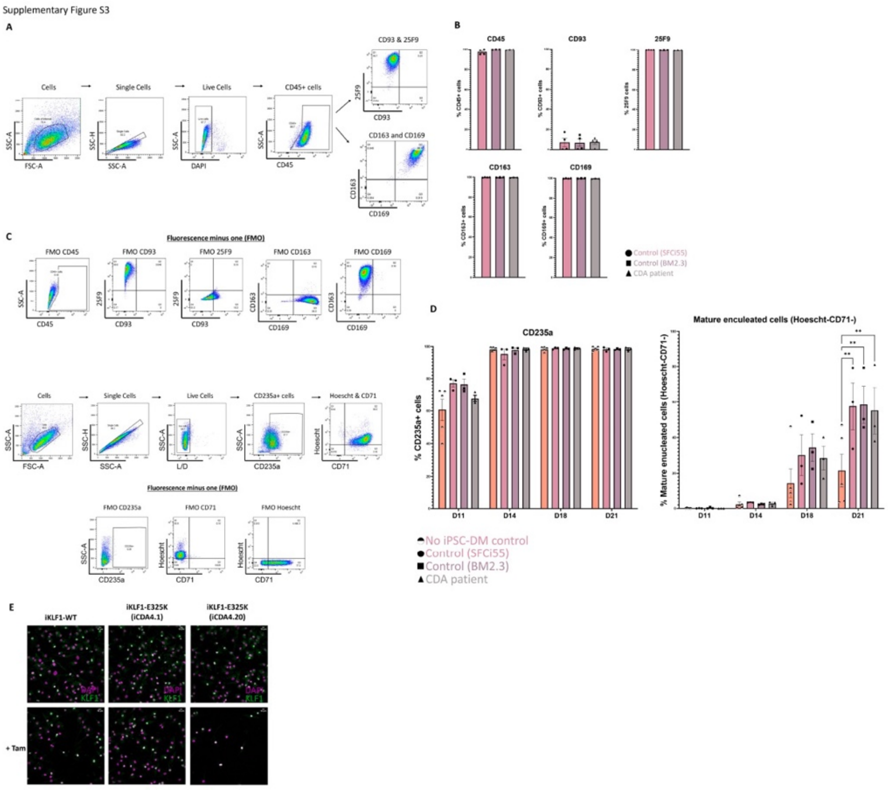


**Supplementary Figure S3:**
A) Gating strategy for analyses of iPSC-DMs. Single, live cells were gated and analysed for expression of CD45, CD93 and 25F9, or CD163 and CD169 using fluorescence minus one (FMO) to set gates. B) Quantification of flow cytometry analyses for cell surface marker expression of CD45, CD93, 25F9, CD163 and CD169 on macrophages generated from two control iPSC lines (SFCi55 and BM2.3) and the patient-derived iPSC line. C) Gating strategy for analyses of suspension cells harvested from *in vitro* EBI assays. Single, live, CD235a$^{+}$ cells were gated and analysed for expression of Hoechst and CD71 using fluorescence minus one (FMO) to set gates. D) Quantification of flow cytometry analyses of suspension cells for CD235a, CD71 and Hoechst at days 11, 14, 18 and 21 of an *in vitro* EBI assay. Error bars represent SEM. One-way ANOVA with Tukey post-test. *$p < 0.05$, **$p < 0.01$. E) Immunofluorescence staining of iPSC-DMs from one inducible KLF1-WT (iKLF1-WT, iKLF1.2) and two inducible KLF1-E325K (iKLF1-E325K, iCDA4.1 and iCDA4.20) iPSC lines stained with an anti-KLF1 antibody (green) and the DAPI nuclear dye (magenta) in the presence (bottom panel) and absence (top panel) of 4OH-tamoxifen. 10uM scale bar. 20X magnification.

**Supplementary Table S1: Primers used for PCR and Sanger sequencing.**

| Primer Name | Sequence |
|---|---|
| KLF1 SDM NEB_FW | CGGGTCAGCTTGTCCGAGCGC |
| KLF1 SDM NEB_RV | CCACTACCGGAAACACACGGG |
| pZDonor_FW | CCGTCGACGCTCTCTAGAGCTAG |
| Int_RV | CCGTCGACGCTCTCTAGAGCTAG |
| 5'aavs1_FW | CGGAACTCTGCCCTCTAACGCTGCCG |
| 5'aavs1_RV | CTGCCAGATCTCTCGAGGCCCTGTGG |
| Ext_RV | ACAGCCCCAGGTGGAGAAACTG |
| KLF1 Sanger sequencing | CACTTTGTTCATCCTAGTCCCA |

**Supplementary Table S2: Primers used for qRT-PCR gene expression analyses**

| Primer Name | Forward Primer Sequence | Reverse Primer Sequence |
|---|---|---|
| GYPA | ATATGCAGCCACTCCTAGAGCTC | CTGGTTCAGAGAAATGATGGGCA |
| TFRC | ATCGGTTGGTGCCACTGAATGG | ACAACAGTGGGCTGGCAGAAAC |
| SLC4A1 | CTGCTGGTGTTTGAGGAAGCCT | CACCAGCAGGATGAGCCAGAAG |
| ICAM4 | GGCACCCATTACACTGATGCTC | AGGCACTTGCATAGGTACGCAG |
| HBA1 | GACCCGGTCAACTTCAAGC | AGAAGCCAGGAACTTGTCCA |
| KLF1 WT | ACACCAAGAGCTCCCACCT | GTAGTGGCGGGTCAGCTC |
| KLF1 WT/E325K | ACACCAAGAGCTCCCACCT | GTAGTGGCGGGTCAGCT |
| IL33 | CCACCAAAAGGCCTTCACT | AAGGCAAAGCACTCCACAGT |
| IGFBP6 | CACAGGATGTGAACCGCAGAGA | CACTGAGTCCAGATGTCTACGG |

**Supplementary Table S3: Antibodies used for flow cytometry analyses**

| Cell Surface Marker | Conjugated Fluorochrome | Manufacturer | Dilution |
|---|---|---|---|
| CD43 | APC | 17-0439-42, Invitrogen | 1:100 |
| EpCAM | PE | 324205, Biolegend | 1:100 |
| CD235a | FITC | 349104, Biolegend | 1:200 |
| CD71 | APC | 17-0719-42, Invitrogen | 1:200 |
| CD45 | APC | 15577936, Ebioscience | 1:100 |
| CD93 | PE | 10804637, Ebioscience | 1:200 |
| 25F9 | E-FLUOR-660 | 15599866, Ebioscience | 1:20 |
| CD163 | PE-CY7 | 333614, Biolegend | 1:25 |
| CD169 | APC | 346007, Biolegend | 1:25 |
